# Long-term impact of fecal transplantation in healthy volunteers

**DOI:** 10.1101/671644

**Authors:** Oleg V Goloshchapov, Evgenii I Olekhnovich, Sergey V Sidorenko, Ivan S Moiseev, Maxim A Kucher, Dmitry E. Fedorov, Alexander V Pavlenko, Alexander I Manolov, Vladimir V Gostev, Vladimir A Veselovsky, Ksenia M Klimina, Elena S Kostryukova, Evgeny A Bakin, Alexander N Shvetcov, Elvira D Gumbatova, Ruslana V Klementeva, Alexander A Shcherbakov, Margarita V Gorchakova, Juan José Egozcue, Vera Pawlowsky-Glahn, Maria A Suvorova, Alexey B Chukhlovin, Vadim M Govorun, Elena N Ilina, Boris V Afanasyev

## Abstract

**Background:** Fecal microbiota transplantation (FMT) is now approved for the treatment of refractory recurrent clostridial colitis, but a number of studies are ongoing in inflammatory bowel diseases, i.e., Crohn’s disease, nonspecific ulcerative colitis, and in other autoimmune conditions. In most cases, the effects of FMT are evaluated on patients with initially altered microbiota. The aim of the present study was to evaluate effects of FMT on the gut microbiota composition in healthy volunteers and to track long-term changes.

**Results:** We have performed a combined analysis of three healthy volunteers before and after FMT with frozen capsules, followed by evaluation of their general condition, adverse clinical effects, changes of basic laboratory parameters, and several immune markers. Intestinal microbiota samples were evaluated by 16S rRNA gene sequencing (16S seq) and shotgun sequencing (or whole-genome sequencing – WGS). The data analysis demonstrated the profound shift towards the donor microbiota taxonomic composition in all volunteers. Following FMT, all the volunteers exhibited gut colonization with donor gut bacteria, and persistence of this effect for almost ~1 year of observation. Transient changes of immune parameters were consistent with suppression of T-cell cytotoxicity. FMT was well tolerated with mild gastrointestinal adverse events and systemic inflammatory response in one volunteer.

**Conclusions:** The FMT procedure leads to significant long-term changes of the gut microbiota in healthy volunteers with the shift towards donor microbiota composition, being relatively safe to the recipients without long-term adverse events.

## Background

Intestinal microbiota is a key player in human body metabolism. Gut microbiota begins to develop from birth, and its composition depends on multiple factors: delivery type, nosocomial microflora at the obstetrics unit, maternal diet, breast feeding etc. [1, 2]. The microbiota serves for a number of vital functions supporting the physiological homeostasis, including synthesis of vitamins and essential aminoacids, short-chain fatty acids (SCFA), e.g., butyrate, propionate, acetate which serve as energy substrates for epithelial cells, as well as inactivation of toxic substances [3]. In cases of negative effects, e.g., antibacterial or cytostatic treatment, profound changes in gut microbiota composition are observed, followed by reduced bacterial diversity, and predominance of pathogenic microorganisms that facilitate damage to a gut epithelium barrier and/or alter immune system response [4].

At present, fecal microbiota transplantation (FMT) from allogeneic donors becomes a popular approach to the microbiota correction. Recently, the FMT procedure was approved by the U.S. Food and Drug Administration for application in the setting of clinical trials in recurrent clostridial colitis (*Clostridium difficile* infection – CDI) [5]. However, several procedure limitations still exist, thus precluding wider implementation of this technology, especially in other clinical settings [6].

Growing interest to this method is determined by a high response rate (>90%) in CDI, including cases with multiple antibiotic resistance [7], positive therapeutic effect in severe cases of ulcerative colitis [8], Crohn’s disease [9], as well as by a relatively simple application. There is also evidence of FMT efficacy in correcting microbiota following antibacterial treatment [10]. Some data provide evidence for the place of FMT in a complex therapy of autoimmune diseases [11], antibiotic-associated diarrhea, and in graft-versus-host disease occurring after hematopoietic stem cell transplantation [12, 13]. The FMT procedure results in reduced prevalence of gut *Enterobacteriaceae* with multiple resistance to beta-lactam antibiotics and carbapenems, vancomycin-resistant *Enterococcus* spp., methicillin-resistant *Staphylococcus aureus* [14], *Klebsiella pneumoniae* and other drug-resistant bacteria [15, 16, 17]. These observations are particularly valuable because of the high mortality caused by antibiotic-resistant pathogens [18].

Gut colonization with donor microbiota is the main suggested mechanism of FMT effects in inflammatory bowel diseases (IBD) [19]. Multiple changes of the gut microbiota composition are induced by FMT [20, 21]. However, there is a lack of data on the exact mechanisms behind FMT efficacy. Kump et al. [22] have shown that the changing taxonomic spectrum approaching a donor microbiota is the main factor determining FMT efficacy in patients with ulcerative colitis. Colonization with donor microbiota may, of course, promote metabolic potential of the gut recipient microflora, thus causing clinical improvement. However, these studies were carried out exclusively with severely ill patients, or in animal models. Despite encouraging results with FMT in different clinical settings, we have not found any data on typical effects of FMT in healthy subjects, with respect to evaluation and tracing the microbiota shifts and appropriate immune system changes, thus allowing to better understand these changes in different clinical disorders. Such data may specify the patterns of the changing host-donor microbiota interrelations after FMT.

The aim of our study was to evaluate effects of FMT upon gut microbiota in healthy persons following FMT from a healthy donor, as well as basic parameters of the immune system before and after this procedure.

## Methods

### Donor selection

A women, 36 years old, was chosen as a single donor of the gut microbiota (body mass = 54 kg, BMI = 19.4) used for all the FMT recipients. She received a usual, balanced, European diet over the entire period of the study and was clinically evaluated according to a protocol recommended by the European Consensus Conference on Faecal Microbiota Transplantation in Clinical Practice [23]. Three donor samples were collected: baseline (was used for FMT) and after 193 and 385 days after baseline

### Selection of the volunteers and treatment schedule

Physically and mentally healthy volunteers participated in this study. They kept a standard European diet (the summary data on the volunteers and the donor are presented in Supplementary Table S1). Detailed information on the aims, tasks and expected results of this study was presented to the volunteers, the FMT procedure was explained in detail. Before entering the study, the volunteers signed an appropriate informed consent.

The FMT procedure was performed in three healthy volunteers (38.6±7.4 years old). Volunteers were assigned IDs - V1 (male), V2 (male) and V3 (female). V2 and V3 recipients are family pair. The treatment was two-staged: the first stage consisted in the administration of 15 capsules containing donor stool on one day, and 15 capsules on the next day. Mild breakfast was allowed 4 hours before administration. One hour before treatment, each volunteer took a dose of omeprazole (20 mg). The volunteers were administered solid gelatin capsules (Coni-Snap^®^ size 4) containing frozen at −20 °C feces, followed by drinking water. The anticipated weight of the material for each single volunteer was 22 g in 30 capsules.

Follow-up continued for 300-303 days. In total, 22 fecal and blood samples of volontaires were collected. The gut metagenomic study was performed for the first volunteer in ten time points; he underwent two FMTs (the second FMT was carried out 38 days after the first FMT). The second volunteer (V2) was administered half of the anticipated dose due to the development of systemic inflammatory response syndrome (SIRS). The third volunteer was administered the full anticipated dose. Adverse effects (AE) were assessed using the Toxicity Scale (Common Terminology Criteria for Adverse Events (CTCAE) Version 5.0 Published: November 27, 2017)

### Sample collection, preparation and sequencing

Collection of the stool samples was performed in sterile plastic containers, both before FMT and at different terms later on. Dynamic monitoring of clinical blood cell counts, biochemical blood analysis, and evaluation of lymphocyte subpopulations were performed in the recipients. Immunophenotyping was performed with a flow cytometer Cytomics FC500 (Beckman Coulter, USA) with CXP Analysis software (Beckman Coulter) using fluorochrome-labeled monoclonal antibodies (CD45 FITC/CD4 PE/CD8 ECD/CD3 PC5, CD19PC7, CD3 FITC/CD(16+56) PE, CD45 PC5, CD5 FITC/CD23 PE/CD19 ECD, CD27PC7, purchased from Beckman Coulter, USA) and Versalyse wash-free lysis (Beckman Coulter, USA).

#### DNA isolation from stool samples

DNA was extracted using PureLink™ Microbiome DNA Purification Kit (Invitrogen™, USA) according to manufacture protocol.

#### 16S rRNA gene library preparation and sequencing

16S rRNA gene library preparation and sequencing were done according to Illumina protocol (16S Metagenomic Sequencing Library Preparation). Briefly, extracted DNA was amplified using standard 16S rRNA gene primers, complementary to V3-V4 region and containing 5’-illumina adapter sequences. In the next step individual amplicons were PCR – indexed and pooled. DNA libraries were sequenced on a MiSeq instrument (Illumina, San Diego, CA, USA) using Miseq reagent kit v3 (Illumina, San Diego, CA, USA).

#### Shotgun library preparation and sequencing

300 ng of DNA were sheared by sonication with the Covaris S220 System (Covaris, Woburn, Massachusetts, USA). The final sizes of fragmented DNA samples were determined on Agilent 2100 Bioanalyzer (Agilent, USA) using the manufacturer guide, and were approximately 400-500 bp long. Paired-end libraries were prepared according to the manufacturer’s recommendations using NEBNext Ultra II DNA Library Prep Kit (New England Biolabs, USA). The libraries were indexed with NEBNext Multiplex Oligos kits for Illumina (96 Index Primers, New England Biolabs, USA). Size distribution for the libraries and their quality were assessed using a high-sensitivity DNA chip (Agilent Technologies). The libraries were subsequently quantified by Quant-iT DNA Assay Kit, High Sensitivity (Thermo Scientific, USA). DNA sequencing was performed on the HiSeq 2500 platform (Illumina, USA) according to the manufacturer’s recommendations, using the following reagent kits: HiSeq Rapid PE Cluster Kit v2, HiSeq Rapid SBS Kit v2 (500 cycles), HiSeq Rapid PE FlowCell v2 and a 2% PhiX spike-in control.

#### Data analysis

The results of 16S rRNA sequencing were independently evaluated by two different computer tools. First tool: metagenomics 16S rRNA Workflow MiSeq Reporter Package, provided together with Illumina sequencing platform with applied the GreeneGenes database [24]. Second tool: DADA2 pipeline [25] and SILVA – 16S genes database [26] was applied to predict the taxonomic annotation using QIIME2 (https://qiime2.org). Due to the compositional type of such data (CoDa), also to WGS data analysis evaluation required CoDa analysis [27, 28] approaches such as Aitchison’s distance [29, 30] with the aid DEICODE [31].

The sequence quality filtration for the WGS metagenomic data was performed by means of the “metaWRAP read_qc” module [32]. To compare taxonomic compositions for metagenomic WGS data, we used MetaPhlAn2 [33, 34]. CoDa approaches (Aitchison distance) and non-metric multidimensional scaling (NMDS) were used for bi-dimensional visualization. A balance dendrogram (CoDa dendrogram) was used for building a model of ecological succession of recipient gut microbiota due to FMT. This dendrogram-like graph shows: (a) the way of grouping parts of the compositional vector; (b) the explanatory role of each sub-composition generated in the partition process; (c) the decomposition of the total variance into balance components associated with each binary partition [27, 28]. Before the analysis, removal of rare taxa and substitution of zeros by Bayesian estimation of (non-zero) proportions were performed [35].

For additional analysis unweighted UniFrac distance and Bray-Curtis dissimilarity was used. Visualization was performed in the R statistical environment vegan package [36] (Euclidean distance and metaMDS function with default parameters) and ggplot2 library (https://ggplot2.tidyverse.org).

#### Detecting donor bacteria in the recipient metagenomes

To trace distinct donor-derived strains in the recipient metagenomic data we used genome-resolved metagenomic (GRM) approaches based on metagenome-assembled genomes (MAGs). To assemble the MAGs, individual samples from donor and each recipient were used separately. The metaWRAP pipeline was used for the MAGs assembly [32] (contain MEGAHIT [20], CONCOCT [37], MetaBat [38], MaxBin2 [39]), with the following parameters of resulting bins: completeness > 70%, contamination < 10%, nucleotide length > 2,000,000 bp. Multiple alignments for 43 marker MAGs segments (amino acid sequences), plotting a phylogenetic tree, and subsequent taxonomic annotation was performed by means of CheckM [40]. The MAGs were clustered by alignments, guided by 100% amino acid (AA) similarity between the studied sequences (*dist.alignment* from ***seqinr*** package for R [41]). The clusters obtained were then additionally compared by their nucleotide similarity using OrthoANI [42] using full MAGs sequences. To follow the dynamics for donor MAGs in metagenomic samples from recipients, the Anvi’o framework [43] and Bowtie2 (100% similarity) [44] functional was applied, suggesting a design of contig database from the donor-derived MAGs, alignment of metagenomic samples, as well as visualization of resulting data. Additionally, for tracking donor-derived bacteria in recipient metagenomes metaSNV (major allele distance was applied as dissimilarity metric) [45] profiling based on mOTUs2 pipeline database [46] was used.

## Results

### Clinical observations

Clinical observations of the volunteers were performed during the first month post-FMT. The AEs were registered for all the subjects 8 to 10 hours after taking the capsules (Supplementary Table S2). There was no emerging AE past the first 24 hours. V1 and V3 exhibited only grade 1 gastrointestinal AEs post-FMT. The second volunteer (V2) developed a SIRS. Dynamic monitoring of clinical blood cell counts, biochemical blood analysis, and evaluation of lymphocyte sub-populations are presented in Figure 2. On day 2 of treatment all laboratory tests were in normal range, except for increased blood neutrophil counts from 59.1% (5.1 × 10^9^/l) to 70.6% (8.9 × 10^9^/l), and from 61.4% (6.3 × 10^9^/l) to 70.7% (6.9 × 10^9^/l), for V1 and V2, respectively. Blood lymphocyte counts showed a decrease from 31.7% to 23.6%, at similar absolute lymphocyte numbers (2.8 × 10^9^/l and 3.0 × 10^9^/l). V3 exhibited a relative decrease of lymphocyte counts, both in percentage (30.1% to 17.8%) and in absolute values (2.8 × 10^9^/l to 1.7 × 10^9^/l).

**Figure 1.**
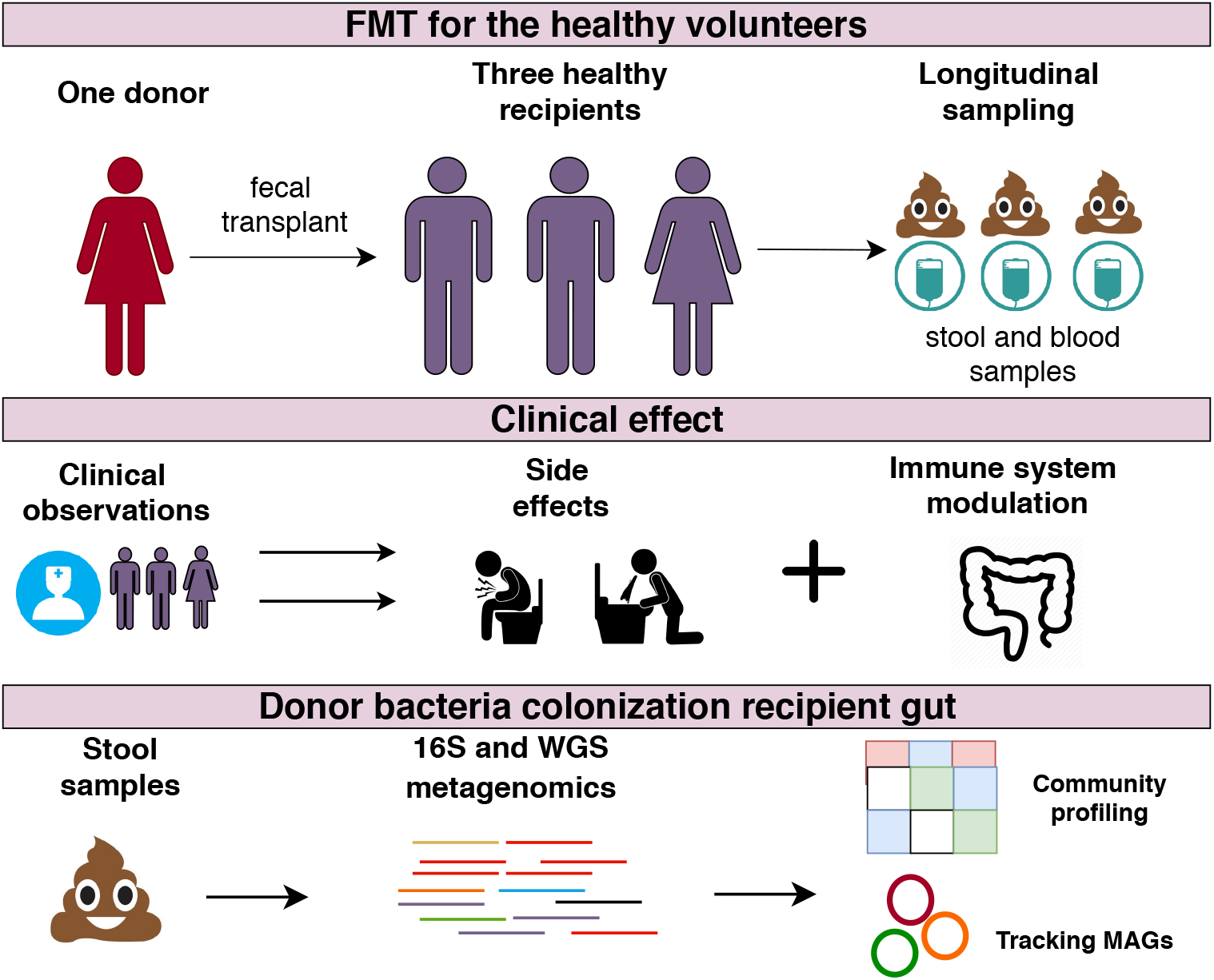
Study description. The first line describes sampling points, the second line clinical effect observable caused by taking FMT capsule. The third line describes obtaining sequencing data and bioinformatic analysis.

**Figure 2.**
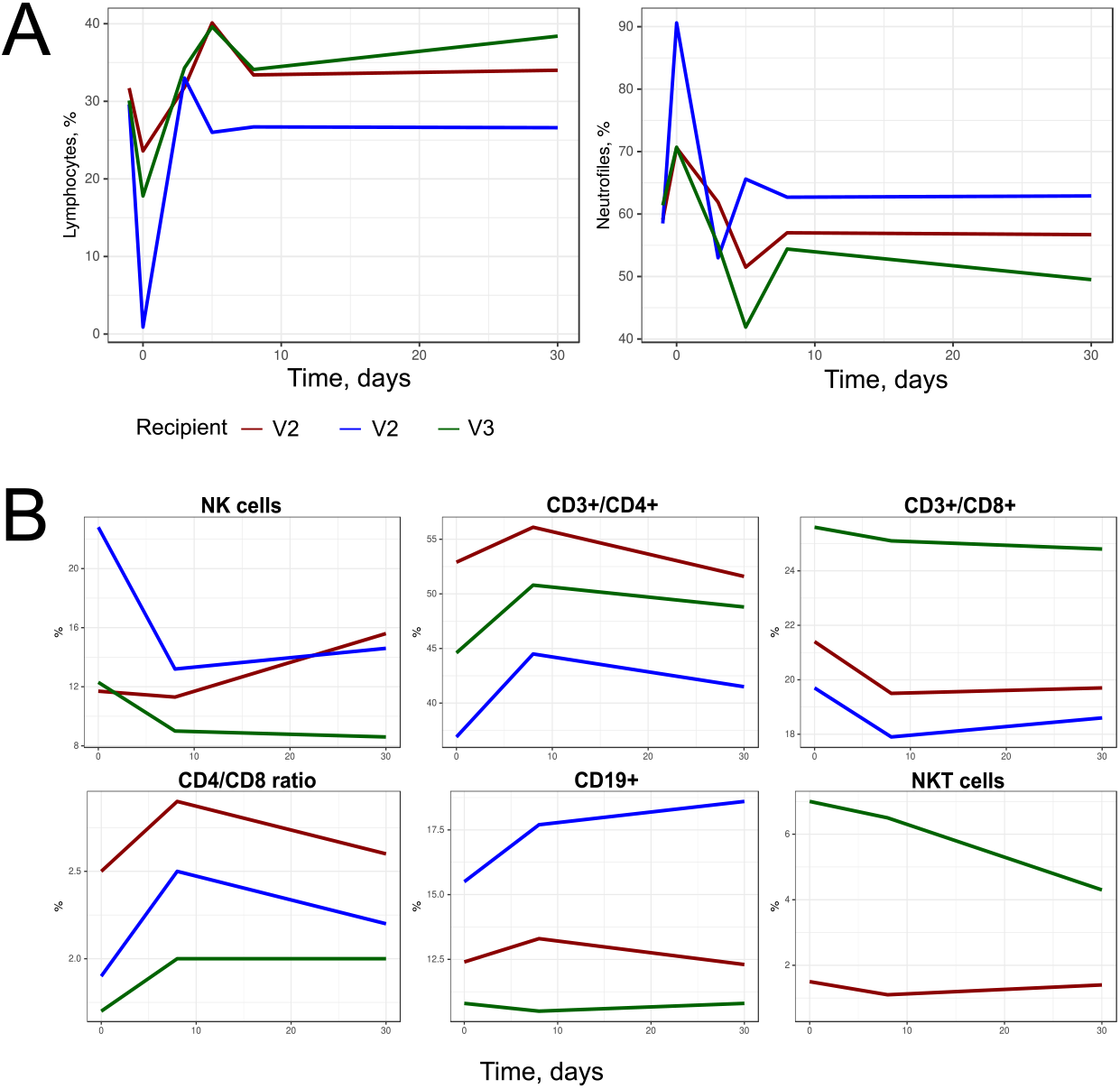
Dynamics of neutrophil counts, lymphocyte counts (A) and lymphocyte subpopulations after FMT (B). The second volunteer (V2) developed SIRS.

For V2, we observed a number of pronounced symptoms, which required additional therapy. Ciprofloxacin was administered at the daily dose of 500 mg for 3 days, and the 2nd round of FMT in this subject was canceled. V2 developed a clinical pattern of systemic inflammatory response (fever, with one-time rise of body temperature to 39.1 °C, with shivers and tachycardia of 102 per minute on the day after administration). The blood changes corresponded to acute bacterial infection: leukocytosis to 16.7 × 10^9^/l, neutrophils 90.6% (15.1 × 10^9^/l), absolute lymphopenia (0.9%, 0.2 × 10^9^/l). Blood smear counts showed increase in band forms, 10% (1.67 × 10^9^/l), segmented forms, 80% (13.36 × 10^9^/l); toxic granulation in the neutrophils and decrease of lymphocytes, 4% (0.66 × 10^9^/l). C-reactive protein levels was within normal ranges, a marginal increase of γ-glutamyl TP to 56.7 U/l (normal values, 0-55 U/l) and ALT to 62 U/l (normal ranges, 0-50 U/l) was noted on day 2. Clinical chemistry parameters of V1 and V3 were within normal ranges during the treatment course.

The lymphocyte subpopulations were examined before FMT, as well as on day 9 and day 30. By day 9, an increased percentage and absolute numbers were observed for T-helpers CD3+CD4+, CD19+CD23+ cells; CD4/CD8 ratio; as well as a decrease in lymphocyte subpopulations, i.e., T-cytotoxic CD3+CD8+ lymphocytes, and NK cells (CD3-CD16+56+). By day 30, a reverse dynamics to normal values was revealed. The number of recipients was insufficient to evaluate the statistical significance of the observed changes.

### Gut microbiome changes after FMT

16S rRNA gene sequencing (16S seq) data analysis was performed in two independent laboratories and the results were consistent in both assays (Supplementary Table S3). The bi-dimensional plot obtained with 16S seq taxonomic data is presented in Figure 3. It shows the convergence of the recipients gut taxonomic profiles to the donor profile within 300 days after the FMT. Interestingly, the gut metagenomic profile of V1 showed a dramatic change after the second FMT procedure from the same donor (2 days after the second FMT). However, further samples showed a return to the donor pattern. Additionally, analysis with using NMDS and unweighted UniFrac distance confirmed previous results (see Supplementary Figure S1A).

**Figure 3.**
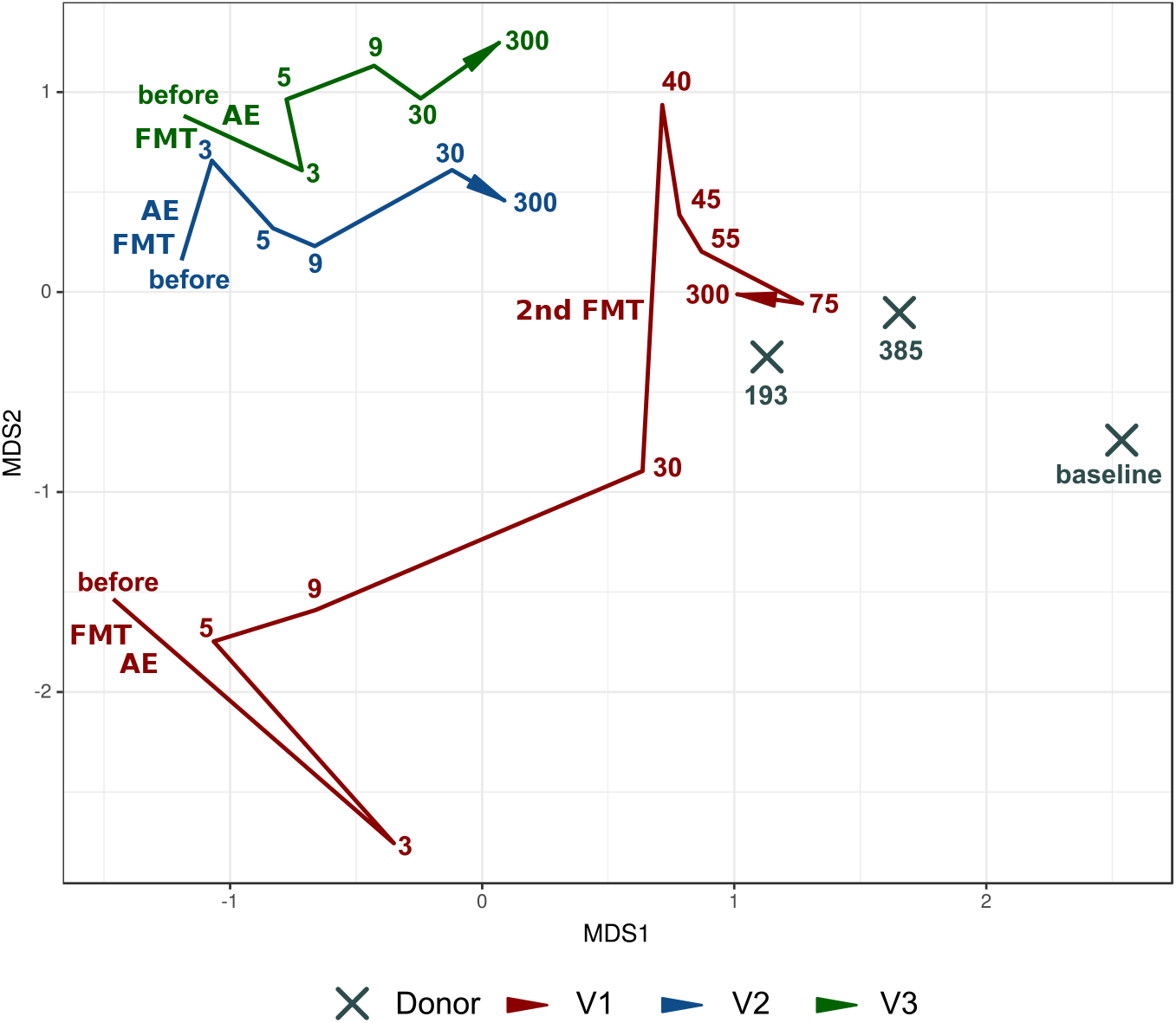
Movement of recipient samples to the donor during the observation time based on 16S rRNA gene sequencing taxonomic composition. Non-metric multidimensional scaling bidimensional obtained Aitchison distance with the aid of DEICODE. Donor samples: X. Volunteer’s samples: red / blue / green colors (see figure legend). The lines denote the evolution of the volunteer’s samples in time (different time points). The days after FMT procedure (or baseline for donor samples) denoted by color numbers. Time of FMT and adverse effects (AE) are schematically noted.

Shotgun metagenomic sequencing was another method for studying changes in the intestinal microbiota profile of the recipients, which yielded 23.1 ±3.7 M of 250 bp reads per sample (98.3 Gbp in total) after quality control. Seventeen metagenomic samples were sequenced with the shotgun method (6 for the V1, 4 for the V2 and V3 and 3 samples for the donor, see sampling scheme in Supplementary Figure S1). The sequencing summary statistic is presented in Supplementary Table S4. A total of 74 genera was detected in all samples. The dataset of relative abundances of bacterial genera is shown in Supplementary Table S5. The shotgun sequencing confirmed the 16S seq data with a similar pattern of changes towards the donor profile (Figure 4A). Similar results was obtained by NMDS bi-dimensional visualization using Bray-Curtis dissimilarity (see Supplementary Figure S1B).

**Figure 4.**
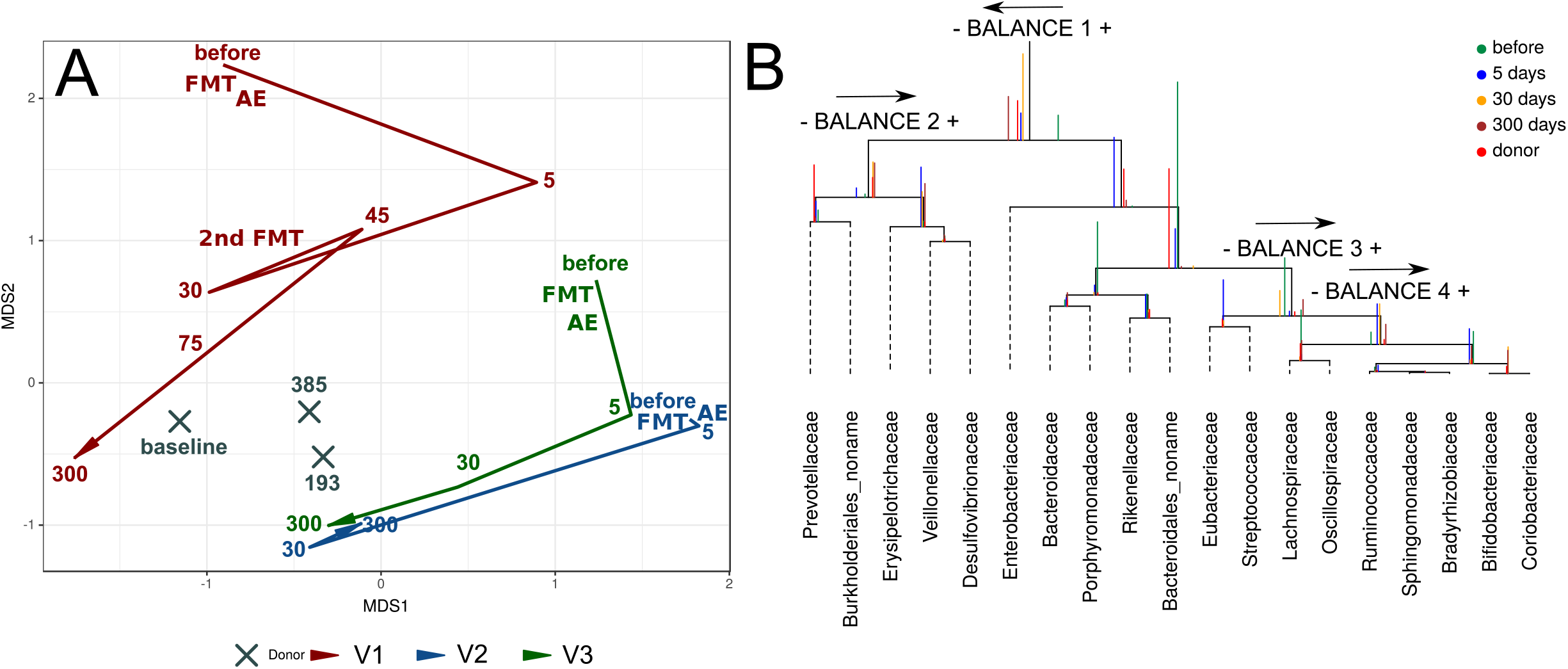
Shifts of the taxonomic profile of microbiota in volunteers towards donor values over the observation time. The figure is based on the shotgun sequencing data. **(A)** Bi-dimensional plot of MetaPhlAn2 taxonomic profile (genera level relative abundances), based on the Aitchison distance. The days after FMT procedure (or baseline for donor samples) denoted by numbers. Time of FMT and adverse effects (AE) are schematically noted. **(B)** CoDa dendrogram which characterizes association of bacterial families, balances presented as edges. Decomposition of total variance by balances between groups of families is shown by vertical bars. Mean values of balances is shown by anchoring points of vertical bars. Color of vertical bars corresponds to time points. Color rectangles highlighted families belonging to important balances. The arrows direction indicates the predominance of this balance part in the donor.

For constructing the model of microbiota succession caused by FMT the balance dendrogram (CoDa dendrogram) was used. This approach allows to identify specific balances (ratio between taxonomic abundances) which are involved in the reshaping of the microbiome of recipients [27, 28]. This model describes intensity of taxonomic reshapes when moving the recipients profiles to the donor-specific parameters (see Figure 4B). Immediately on the fifth day after FMT, the recipients relatively increased the content of *Prevotellaceae*, unknown *Burkholderiales, Erysipelotrichaceae, Vellonellaceae* and *Desulfovibrionaceae*; however the shift towards *Prevotellaceae*, unknown *Burkholderiales*, was more pronounced at day 5. At the same time, the relative increase of *Lachnospiraceae*, *Oscillospiraceae*, *Rum-minococaceae, Sphingomonadaceae, Bradyrhizobiaceae, Bifidobacteriaceae* and *Coriobacteriaceae* occurred less quickly and more smoothly. Also, relative abundance of *Enterobacteriaceae*, *Bacteroidaceae*, *Porphyromonadaceae*, *Rikenellaceae*, unknown *Bacteroidales*, *Eubacteriaceae* and *Streptococacceae* decreased gradually towards the donor-like profile.

It is worth to note that the results obtained show a directed change in the gut microbiota composition of volunteers. Thus, the pre-FMT profiles of the recipient microbiota become similar to the donor microbiota post-FMT, as well as to one another.

### Identification of donor bacteria in the recipient metagenomes

Taxonomic profiling methods may reveal general changes of the taxonomic profile for the gut microbiota. It is however important to examine engraftment of the donor bacteria in recipients. To assess engraftment of the donor bacteria using the obtained shotgun sequencing data, we used an approach allowing to restore bacterial genome from the metagenomic data (metagenome-assembled genomes – MAGs). This method is based on the metagenomic assembly and clustering of contigs through a binning procedure. As a result, 243 MAGs were assembled for all metagenomic samples both from donor and recipients. For the donor 46 MAGs were obtained, for each of the volunteers 87, 56, and 54 MAGs, respectively (note that these MAGs represent microbes from both pre- and post-FMT time points). Further, based on 43 marker single-copy proteins, the place at the dendrogram for each MAG was determined (see Figure 2), and appropriate taxonomic annotation was ascribed with the CheckM tool. We detected 14 donor-like MAGs in which 100% amino acid similarity of marker proteins was observed (see Figure 5A). The changes in relative abundances of these 14 MAGs are shown in Supplementary Figure S3. The similarity of the nucleotide sequence between donor and recipient MAGs was also high (see Figure 5B). Anvio visualization for the mapping results of the reads from recipient samples in the donor MAGs is shown in Figure 5C.

**Figure 5.**
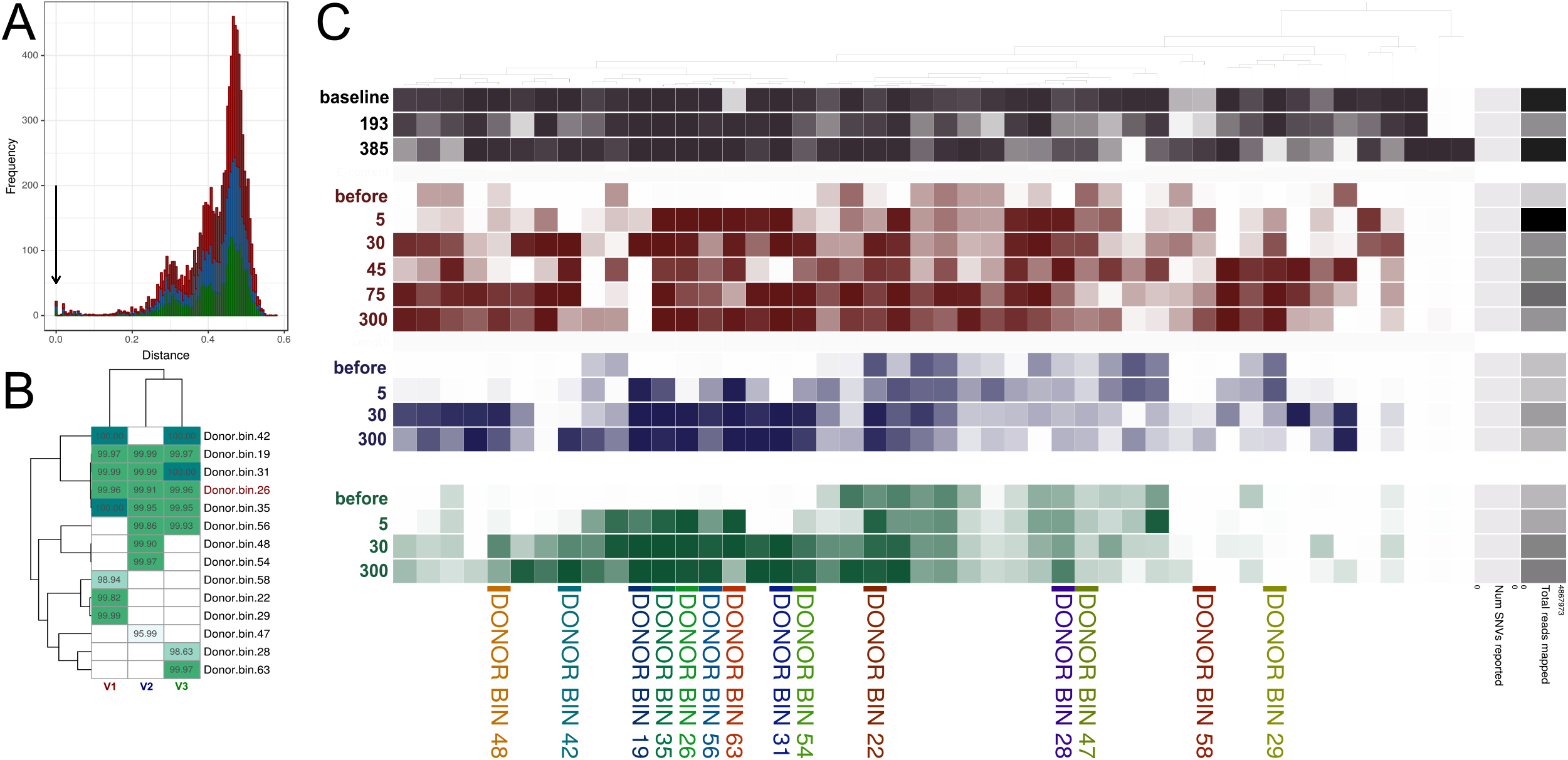
Comparison of similarity between donor and recipients metagenome-assembled genomes (MAGs). **(A)** The AA distance based on 43 marker proteins between all donor MAGs and all MAGs of all recipients. Arrow shows that some MAGs in donor and recipient is present with absolute similarity of marker genes sequence. **(B)** The average nucleotide identity (ANI) between similar donor and recipients MAGs. The MAGs with 100% AA similarity of 43 marker proteins were selected. **(C)** Anvio plot denoted prevalence of donor MAGs across all metagenomic samples. Detection value (proportion of nucleotides in a contig that are covered at least 1x (according to http://merenlab.org/2017/05/08/anvio-views) was used as an abundance metric, which is shown as color brightness. Black color denotes detection value of donor MAGs in the donor samples, red – in the V1 samples, blue – in the V2 samples, green – in the V3 samples. DONOR BIN – clusters of metagenome-assembled genomes similar to the donor bacteria. The days after FMT denoted by numbers. The mapping of recipient metagenomic reads to donor MAGs was performed with 100% similarity.

DONOR_BIN_26 didn’t show the 100% amino acid homology with MAG of V1 and V3. However, they were similar in their nucleotide composition. These discordances could be explained by some metagenomic assembly artifacts and binning, thus resulting in chimeric contigs. The given approach shows a rather big number of false-negative results; however, it allows to detect successful cases from nonspecific findings.

Of 14 donor MAGs with complete amino acid sequence similarity in marker proteins, 10 may be considered as successfully engrafted in at least one recipient. The DONOR_BIN_28, DONOR_BIN_47, DONOR_BIN_22, DONOR_BIN_22, DONOR_BIN_22 did not enter this list due to the following reasons: (1) nucleotide identity from recipient MAGs (threshold <99.90% nucleotide identity); (2) they were covered by reads after FMT (not 100% certainty of their donor origin). By taxonomic annotation, these “strong” colonizers belong to the following orders: *Bacteroidales* (n=5), *Clostridiales* (n=3), *Selenomonadales* (n=1). Interestingly, many donor MAGs didn’t show 100% similarity in the two parameters described above. However, they appeared in the recipient after FMT. This can be explained by the chimeric contigs when assembling recipients MAGs. In addition, similar recipient bacteria can increase after FMT.

Ten MAG clusters, similar in amino acid sequences of marker proteins, were exclusively present in the recipient metagenomes (see Supplementary Figure S3). Based on the criteria of a nucleotide similarity (ANI > 99.90%), there is evidence that the observed changes of several genera of the bacteria abundances were donor-independent. Either these expanding bacteria were in donor samples, but were not found due to insufficient read coverage, or the FMT procedure induces the expansion of certain types of recipient bacteria (for example, see Supplementary Figure S3A). We have also revealed 4 cases of similar MAG sets in V2 and V3 that decreased relatively after FMT in both recipients. V2 and V3 have similar patterns of decrease and increase of some similar MAGs (see Supplementary Figures S3 D, F, H, I, J and Supplementary Figure S4). A more detailed information about MAGs assembly is presented in Supplementary Table S6.

Additionally, SNV-profiling based on mOTUs (phylogenetic marker gene based operational taxonomic units) was performed (see Materials and methods section). After FMT number of mOTUs identical to donor were increased. The results are present in Supplementary Figure S5.

## Discussion

FMT is now increasingly used for treatment of different disorders. Most studies are concentrated on evaluating FMT consequences in patients with CDI, ulcerative colitis, and Crohn’s disease. It was speculated that in these diseases the therapeutic effect is based on the expansion of the donor flora and on the correction of defects in the species composition [47, 48, 49]. The correction of the intestinal microbiome leads to the restoration of short chain fatty acids and bile acid metabolism, altered immune response, the profile of cytokines and chemokines, and augmentation of intestinal wall reparation. The mentioned processes may be immediately or indirectly affected by other medications, e.g. granulocytic colony-forming factor, glucocorticosteroids used in hematopoietic stem cell transplantation, antibiotics used in pseudomembranous colitis, aminosalicylates, anti-TNF monoclonal antibodies in Crohn’s disease, and nonspecific ulcerative colitis. While change in microbiome after FMT is an established fact, it is unknown whether it is the primary therapeutic agent. The study of the FMT effects in healthy volunteers is important to understand the mechanisms behind the efficacy of this procedure and potentially to improve its outcome for patients (e.g. by rules of donor selection).

The present study resulted in several important findings. First, it demonstrated that FMT even with a small stool (bacterial) mass (11-22 g) induces profound changes in healthy persons with normal microbiota composition. The convergence of the recipient taxonomic composition to the donor-like state following the FMT procedure was demonstrated for various diseases [50, 51, 49, 52], but comparable changes were observed in the healthy volunteers. Thus, the effects of FMT are comparable in the normal and pathological conditions, indicating that the replacement of the missing bacterial populations is not a unique. Second, we observed that the composition of the gut flora is altered due to engraftment of donor bacteria. Although this might be a sequencing artifact, all published studies with FMT indicate significant increase in the overall bacterial diversity which is not only related to the growth of the donor flora [53, 54]. The potential mechanisms behind the activation of recipient flora might be horizontal gene transfer [55], effects of the non-bacterial stool components [56], and functional interactions between microbial communities [57]. Further studies are required to elucidate the exact mechanism(s). However, we have identified some features of the restructuring process. *Paraprevotelaceae* and unknown *Bulkholderiales* colonize faster than others. This might be a “hub” bacteria which allows to develop a new “version” of the recipient gut community, which included recipient-derived and donor-derived characteristics.

The number of potential applications of FMT is exponentially growing: decolonization from antibiotic-resistant bacteria prior to stem cell transplantation [54], modulation of response to cancer immunotherapies, like anti-PD-1 antibodies [58], vaccination against respiratory pathogens [56], amelioration or prevention of non-gastrointestinal infections, like malaria [59], treatment of autism [60] and depression [61]. However, there was no evidence that FMT induces long term changes in subjects with no previous damage to microbiota due to antibiotics treatment or to an underlying condition. This study provides the first proof of principle that, even in a healthy person, the procedure induces long term changes with a shift towards donor profile. The FMT in healthy recipients in this study induced several gastrointestinal adverse events and inflammatory response in form of a shift to band leukocyte forms, however with normal CRP levels, thus suggesting massive antigenic exposure, but no bacteremia with development of septic state. A recent detailed review on the side effects in FMT was based on data obtained from 1998 to 2015 [62]. Severe AEs were mostly related to endoscopic procedures and aspiration. The use of capsule FMT seems to minimize the risk of the procedure. The observed mild adverse effects may require correction using only symptomatic therapy, like antiinflammatory and spasmolytic drugs. Leukocytosis and neutrophilia, along with relative and absolute lymphopenia may be a near-normal variant following FMT. One volunteer did receive a short treatment of ciprofloxacin for systemic inflammatory response syndrome (SIRS), but the sequencing data still indicated the shift towards the donor pattern of microbiota. Although early antibiotic use was reported to compromise the efficacy of the procedure [63], in our study the short-term usage of antibiotic without broad spectrum activity had no long-term impact.

Although the study group was small, the observed changes were consistent with previous preclinical studies [64, 65] and case reports [66]. There was transient decrease in total lymphocyte count, decrease of CD8+ cells, decrease of NK cells and increased CD4/CD8 ratio. The effect was most prominent in the volunteer with SIRS. The downregulation of lymphocyte response might have comparable mechanisms to the one induced under bacterial septic conditions [67]. This AE might be one of the mechanisms behind the FMT benefit in the autoimmune disorders.

In the clinical studies it was demonstrated that the efficacy of FMT was based on such genera as *Ruminococcaceae, Lachnospiraceae* and *Prevotellaceae* [53, 60]. These were the bacterial species that were consistently expanded in the volunteer samples. It is unclear if they directly drive the therapeutic effect of the FMT or are just a marker of the changes that induce the response. The question regarding the optimal donor for each indication is still open. Only accumulation of clinical observations will give the answer to this intriguing question. Another approach used by the industry in the early clinical trials is the mixture of products from a large number of donors (https://clinicaltrials.gov/ct2/show/NCT03497806), but the benefits and drawbacks of such an approach are still to be evaluated.

## Conclusions

The main conclusion of the present study is the confirmation of the long-term microbiota composition conversion by FMT in healthy subjects. The microbiota taxonomic composition in recipients shifted towards the donor profile. The most important finding was the expansion of donor-derived bacteria inside healthy recipients gut. Additional important findings may be certain rules of community succession after FMT. Perhaps in the future, the description of these rules will allow the microbiota to be controlled and directed from one state to another.

## List of Abbreviations

FMT: Fecal Microbiota Transplantation
OTU: operational taxonomic unit
MAG: Metagenome-assembled genomes
NMDS: Non-Metric Multidimensional Scaling
CDI: Clostridium Difficile Infection
CoDa: Compositional Data
WGS: Whole Genome Sequencing
ANI: Average Nucleotide Identity
GRM: Genome-Resolved Metagenomics
SNV: single nucleotide variants

## 1 Ethics approval and consent to participate

The study was approved by the Pavlov University Review Board (Protocol No.192, Jan 30, 2017). Before the start of the study, each patient signed an informed consent.

## 2 Availability of data and materials

We made data obtained in this study publicly available at (download.ripcm.com/add_files), where are presented 1) the QIIME2 and MetaPhlAn2 taxonomic tables; 2) the Anvi’o profile to interactively visualize and further investigate donor MAGs mapping profiles across samples; 3) the distribution statistics for each donor MAG across samples; 4) FASTA files for each MAGs. Raw metagenomic data are also deposited at the NCBI Sequence Read Archives under the BioProjects accession numbers PRJNA509769 and PRJNA510036.

## Author’s contributions

GOV – research idea, performed general coordination. OEI – 16S rRNA gene and shotgun sequencing data analysis, partially wrote the manuscript. SSV – 16S rRNA sequencing, edited the manuscript. MIS – general statistical evaluation, edited the manuscript. KMA – wrote the manuscript. FDE – implementation of SNV-based profiling. PAV, MAI – contributed to the bioinformatics analysis and manuscript preparation. GVV – performed 16S rRNA gene sequencing. VVA and KKM – performed shotgun sequencing. KES – supervising of shotgun sequencing. BEA – 16S rRNA gene sequencing data analysis. SAN, GED, KRV, SAA – contributed to the organization of work, edited the manuscript. GMV – performed Immunophenotyping of peripheral blood lymphocytes. EJJ and P-GV – performed CoDa analysis and text manuscript preparation. SMA – preparation of capsules with fecal microbiota. CAB – coordinated routine infectious diagnostics, edited the manuscript. IEN – supervised WGS sequencing and data analysis. GVM and ABV – general management.

## Acknowledgements

We thank the hospital and laboratory staff, volunteers for participating in this study.

## Supplementary Files

Supplementary Figure S1 – Non-metric multidimensional scaling bi-dimensional plots of MetaPhlAn2 taxonomic profile (genera level relative abundances), based on the unweighted UniFrac distance (A) and Bray-Curtis dissimilarity (B). The lines denote the evolution of the volunteer’s samples in time (different time points). The days after FMT procedure (or baseline for donor samples) denoted by numbers. Time of FMT and adverse effects (AE) are schematically noted.

Supplementary Figure S2 – Recipient MAGs with donor MAGs 100% amino acid similarity of 43 marker proteins relative abundance change.

Supplementary Figure S3 – Similar recipient MAGs (with 100% similarity of 43 marker proteins) relative abundance change.

Supplementary Figure S4 – The ANI between similar recipient MAGs. Recipient MAGs with 100% AA similarity of 43 marker proteins were selected.

Supplementary Table S1 – Summary data about the donor and recipients.

Supplementary Table S2 – Adverse effects after FMT in healthy volunteers (scored by Common Terminology Criteria for Adverse Events (CTCAE) Version 5.0.

Supplementary Table S3 – Sequencing general statistics (16S rRNA seq).

Supplementary Table S4 – Sequencing general statistics (shotgun seq).

Supplementary Table S5 – Relative abundance of microbial genera and viruses in the WGS metagenomes (the percentage of overall abundance; obtained using MetaPhlAn2).

Supplementary Table S6 – MAGs assembly and taxonomic annotation statistic.

