## Supplementary figures and images for "Long-term impact of fecal transplantation in healthy volunteers"

### Supplementary Figure 1

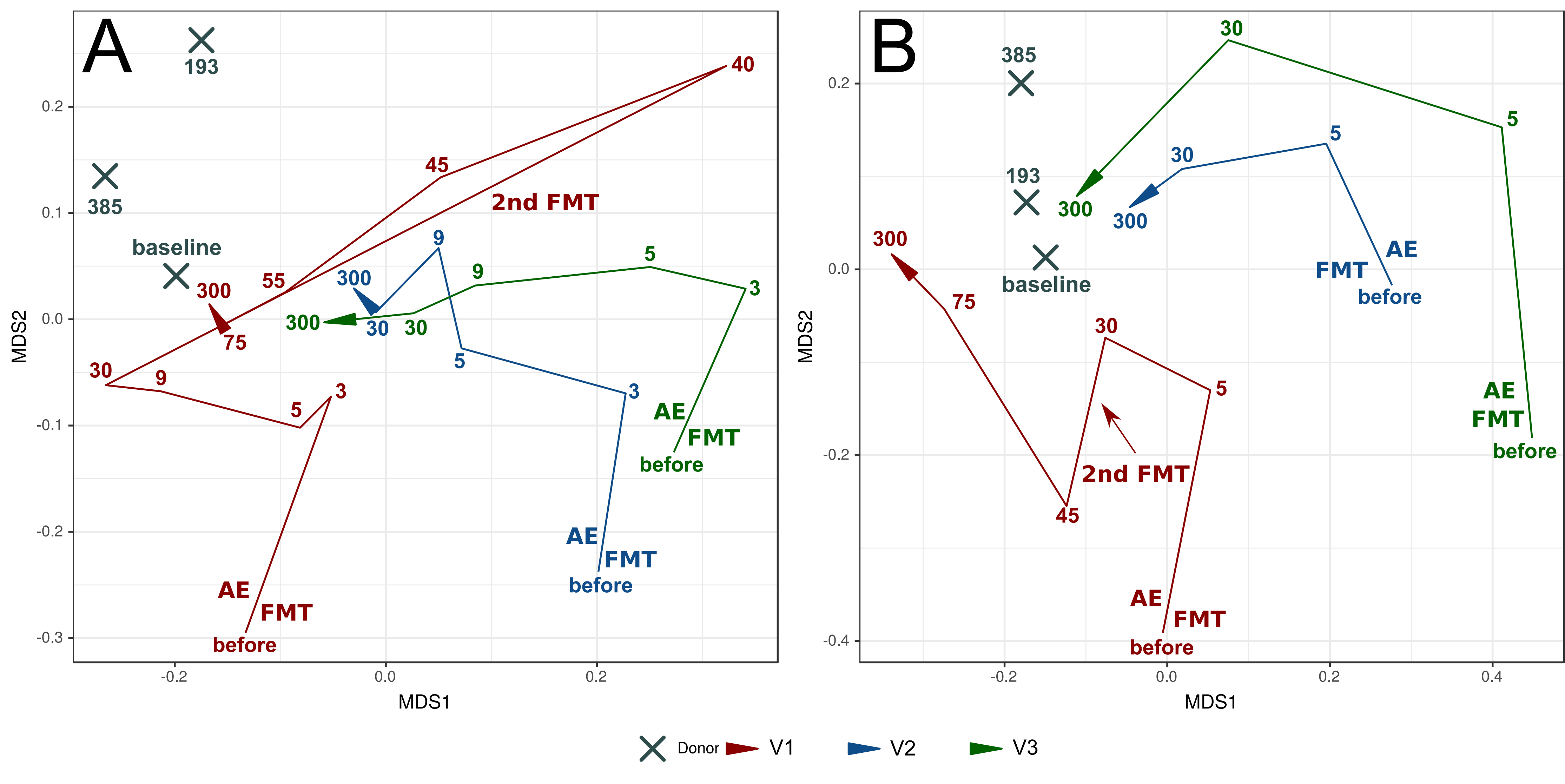

### Supplementary Figure 2

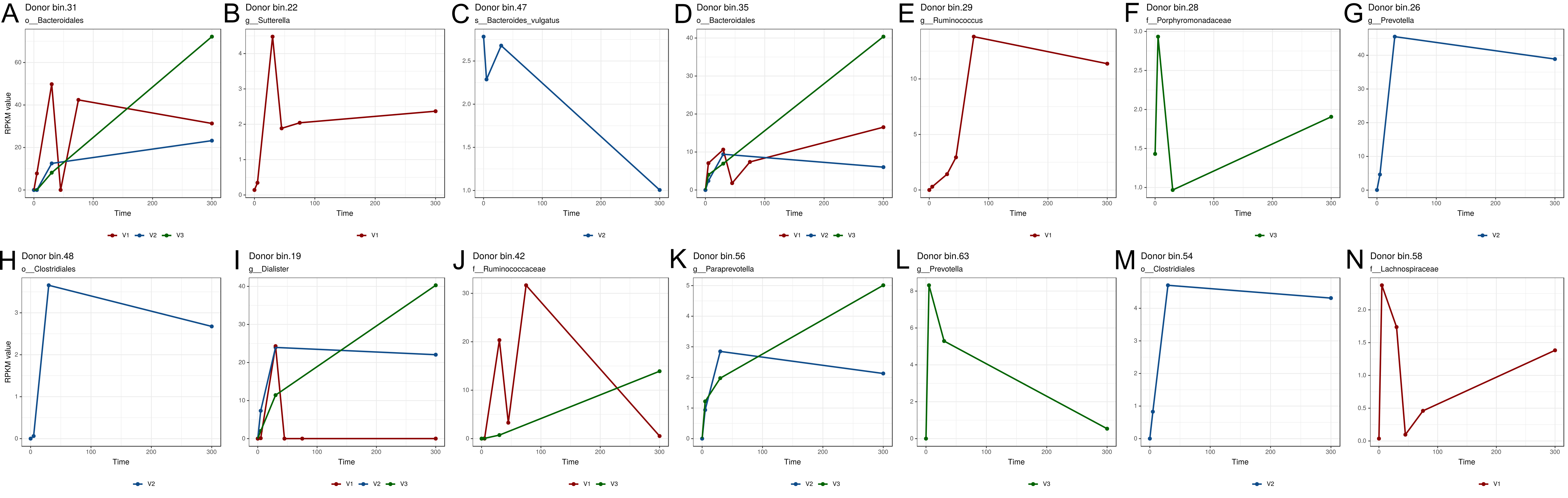

### Supplementary Figure 3

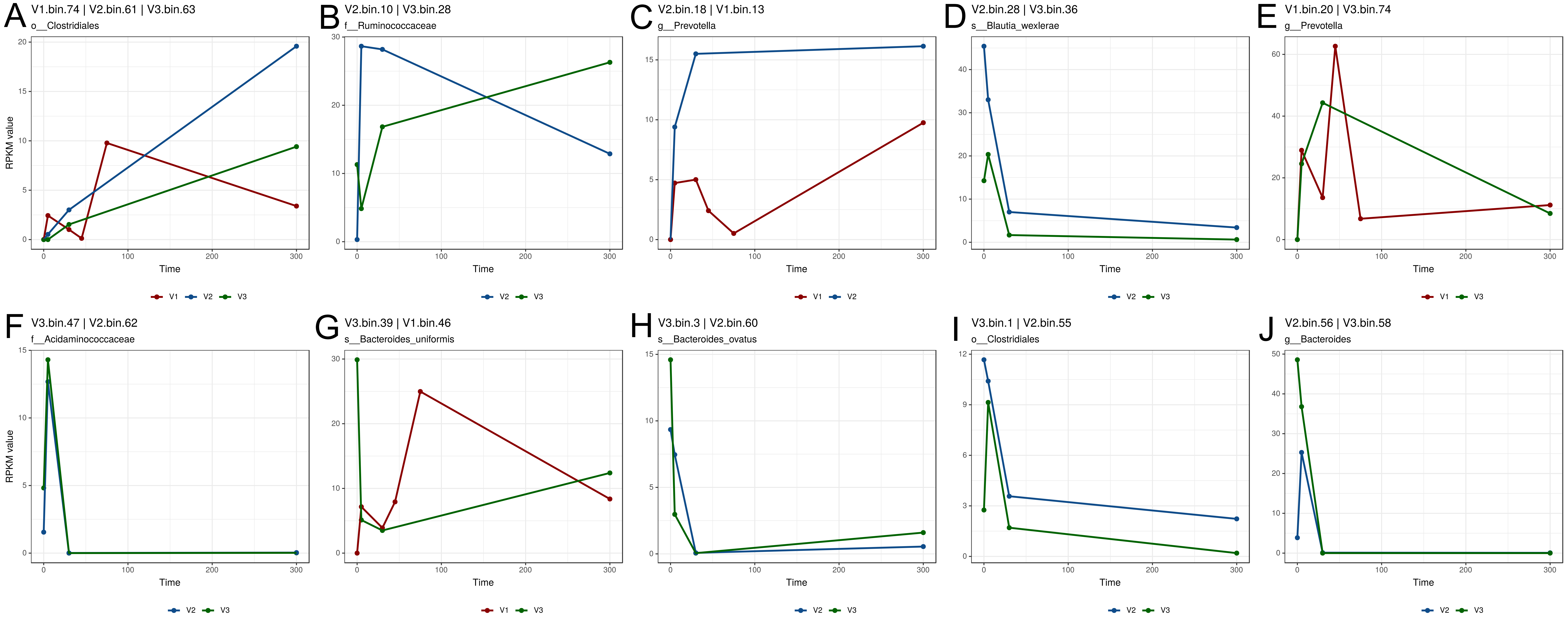

### Supplementary Figure 4

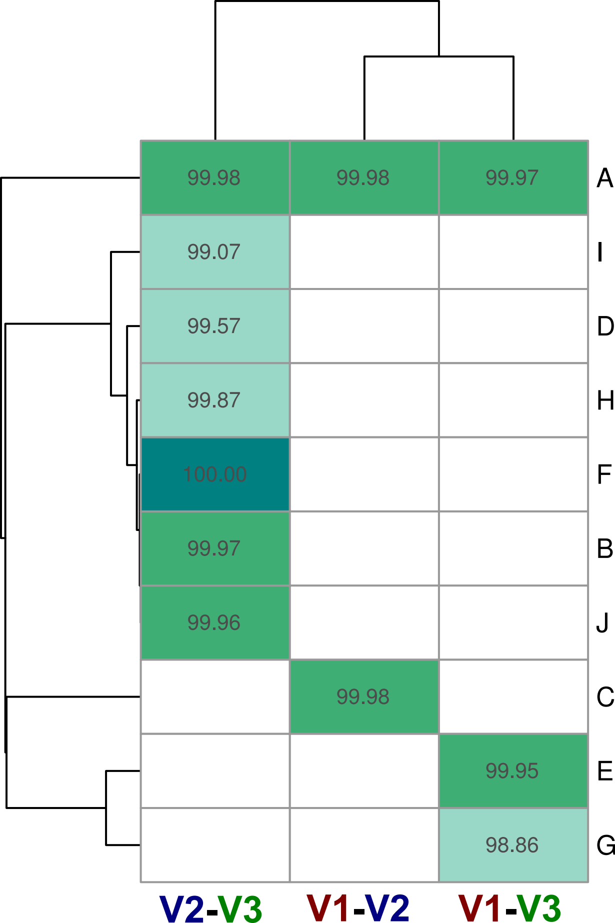

### Supplementary Figure 5

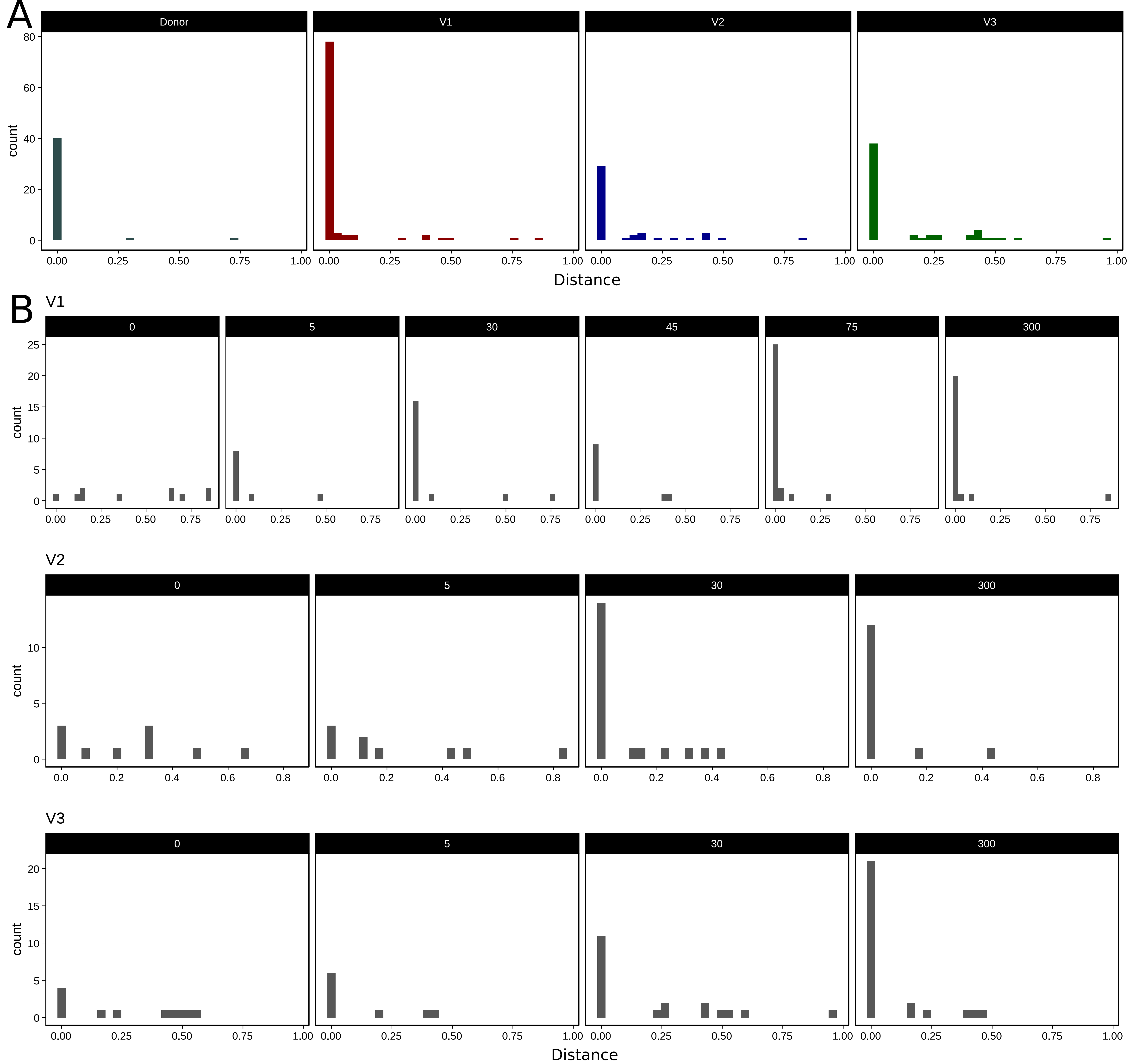
